# Reconstructing the Single-Cell Spatiotemporal Dynamics of Glioblastoma Invasion

**DOI:** 10.1101/2025.03.20.644331

**Authors:** Hitesh Bhagavanbhai Mangukiya, Madeleine Skeppås, Soumi Kundu, Maria Berglund, Adam A. Malik, Cecilia Krona, Sven Nelander

**Author notes:** Correspondence: Sven Nelander, PhD, Department of immunology, genetics and pathology, Dag Hammarskjölds väg 20, 751 85 Uppsala, Sweden. These authors contributed equally to this work: Hitesh Bhagavanbhai Mangukiya, Madeleine Skeppås.

## Abstract

Glioblastoma invasion into healthy brain tissue remains a major barrier to effective treatment, yet current models fail to capture its full complexity in a scalable and patient-specific manner. Here, we introduce GlioTrace, a novel *ex vivo* imaging and AI-based analytical framework that enables real-time, spatiotemporal tracking of glioblastoma invasion dynamics in patient-derived glioma cell culture xenograft (PDCX) brain slices. By integrating whole-specimen confocal microscopy, vascular counterstaining, and an advanced computational pipeline combining convolutional neural networks and Hidden Markov Models, GlioTrace identifies distinct invasion modes, including dynamic morphological switching, vessel-guided migration, and immune cell interactions and quantifies patient-specific variations in invasion plasticity. Using GlioTrace, we demonstrate that targeted therapies can selectively modulate invasion phenotypes, revealing spatially and temporally distinct drug responses. This scalable platform provides an unprecedented window into glioblastoma progression and treatment response, offering a powerful tool for precision oncology and anti-invasion therapeutic development.

## Introduction

Glioblastoma (GBM) is one of the most aggressive and treatment-resistant cancers [1], marked by rapid growth and widespread infiltration into surrounding brain tissue. Standard therapies, such as the Stupp protocol [2], primarily target the tumor core, but fail to address invasive tumor cells that spread beyond resectable regions. These infiltrative cells are a major driver of treatment failure and disease progression. Extensive dissemination—especially into the brainstem—is linked to poorer survival outcomes [3–7]. Conversely, greater surgical removal of invasive cells correlates with longer survival [8, 9]. Understanding and targeting these invasive properties is therefore a critical priority.

GBM invasion is shaped by a complex interplay of cellular and microenvironmental factors. First, as GBM cells invade the brain at the tumor periphery, they engage in intricate interactions with normal brain structures [10], including co-opting blood vessels as migratory scaffolds [11–13], penetrating white matter tracts [14], and even forming synaptic contacts with neurons [15–17]. Second, GBM cells exist in multiple transcriptional states that resemble normal brain cell types and their precursors [18–22]. These states are differentially distributed across the tumor, raising the possibility that distinct cellular states exhibit unique invasion behaviors. Third, GBM cells demonstrate remarkable plasticity, dynamically altering their morphology and differentiation over time. According to the early ‘go-or-grow’ model, GBM cells switch between migratory and proliferative states [23, 24], though more recent lineage-tracing studies suggest a more intricate hierarchy of cell state transitions, with stem-like cells at the apex [19, 22, 25]. Collectively, these findings highlight that GBM invasion is a dynamic and multifaceted process, yet many aspects remain poorly understood. Can, for instance, plastic transitions or cell-cell interactions be modulated by drugs to target invasion? To what degree do these phenomena vary between patients? To address these questions, a critical challenge is the ability to perform long-term, single-cell observations in a model that captures diverse invasion modes at scale. Such observations could enable the development of data-driven models to characterize GBM morphological plasticity and invasion dynamics.

To bridge this gap, we introduce GlioTrace, a scalable platform for real-time, high-throughput analysis of GBM invasion using patient-derived glioma cell culture xenograft (PDCX) brain slices. By combining whole-specimen confocal imaging with AI-driven computational modeling, GlioTrace enables continuous tracking of thousands of tumor cells in their native microenvironment. Our approach integrates vascular counterstaining, single-cell tracking, and machine learning algorithms—including convolutional neural networks (CNNs) and Hidden Markov Models (HMMs)—to classify invasion phenotypes, quantify phenotypic transitions, and assess patient-specific invasion patterns. Compared to intravital imaging in living animals, GlioTrace dramatically increases observational scale and duration, providing a powerful framework for studying GBM invasion heterogeneity and therapy response. Using this system, we identify distinct GBM invasion modes, reveal dynamic morphological transitions, and demonstrate that targeted therapies can modulate invasion phenotypes in a spatially and temporally distinct manner.

Applying GlioTrace to a set of patient-derived models, we uncover key distinctions in GBM invasion strategies. Perivascular migration is dominated by highly motile, translocating cells that engage dynamically with the vasculature, while diffusely invading cells display a more stochastic migration pattern. Notably, we identify a novel class of branching cells exhibiting the highest motility, often serving as intermediates between invasive phenotypes. Using Hidden Markov Models, we establish a hierarchy of morphological states, revealing frequent transitions between translocation and locomotion modes—suggesting a potential regulatory node in GBM invasion. By quantifying invasion dynamics across patient-derived models and testing pharmacological interventions, we demonstrate that invasion is not a uniform process but varies across tumors, with distinct vulnerabilities to anti-invasive therapies. GlioTrace can be implemented on standard high-content microscopes and its software for processing the data and building models is made available as free software free-of-charge. We expect the method to deepen our understanding of GBM invasion and expedite the discovery of tailored anti-invasion therapies for GBM.

## Results

### Whole-specimen monitoring of viable glioblastoma brain slices linked to automated analysis

GlioTrace integrates two components **(Figure 1)**: The first component is an implementation of robust long-term whole-specimen imaging of 300-micron-thick brain slices from established PDCX tumors grown in mouse hosts **(Figure 1A).** In contrast to conventional techniques [10, 24, 26–31], which involve grafting tumor cells directly onto brain slices, we choose to image slices of brains from mice with established, orthotopically grafted, tumors. This approach helps ensure that the tumor is well integrated into the brain anatomy, preserving natural tumor-stroma interactions. The tumor cells are marked by Green Fluorescent Protein (GFP). To relate the tumor cells to the overall brain anatomy and the local tissue environment we apply a tissue-penetrant conjugate between tomato lectin (TL) and a red fluorophore. The conjugate, TL-DyLight 594, is selective for endothelial cells and microglia/macrophages and saturates the specimen within 18 hours **(Supplementary Fig. 1A-B; Supplementary video 1A)**. The incubation protocol was optimized to ensure minimal signs of cellular damage, as detected by ethidium homodimer-1 **(Supplementary Fig. 1C-D).** Over a period of 4-6 days, GlioTrace captures the whole specimen by pooling multiple microscopic fields and Z stacks resulting in data in 10X resolution, sufficient to capture slender cellular protrusions in videos with 9000×9000 pixel resolution, over circa 100 time points (1-2 hour intervals). The second component of GlioTrace is a set of algorithms that stabilize and denoise the data, identify, classify and track cells, and estimate both patient differences and treatment effects **(Figure 1B)**. The GlioTrace software (MATLAB-based) integrates subpixel video stabilization techniques (to adjust for specimen drift), Kalman filtering (to track cells), convolutional neural networks (to classify cell morphologies), Hidden Markov Models (to model cell state transitions), and linear mixed effects models (to summarize treatment effects). Our system thereby enables both population-level analyses and detailed cell tracking, allowing us to capture invasion dynamics in real time **(Figure 1B).** The approach is scalable, capable of analyzing six whole brain PDCX slices in parallel over five days, covering thousands of cells, making it suitable for high-throughput drug screening.

**Figure 1.**
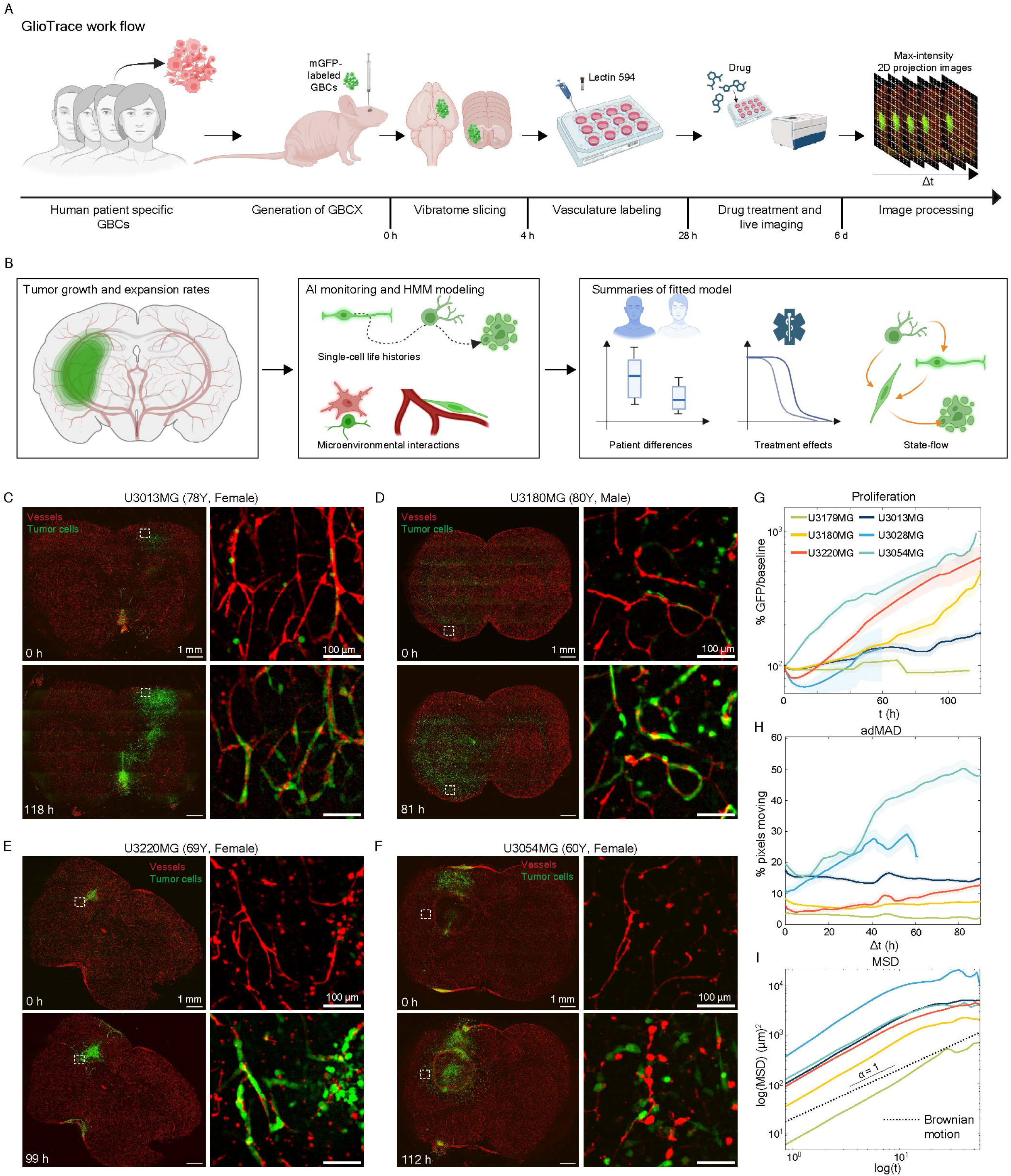
Overview and characterization of glioblastoma PDCX tumors in slice culture. **(A)** Experimental workflow for patient-derived glioblastoma xenograft (PDCX) brain slice culture. **(B)** GlioTrace framework integrating imaging, cell tracking, and computational modeling. **(C)** Representative confocal images showing perivascular migration of bulky glioblastoma cells at the first and last time points. (**D)** Representative images of diffusely growing glioblastoma cells showing both perivascular and diffuse migration. (**E, F)** Additional examples of perivascular migration in bulky glioblastoma tumors. **(G)** Quantification of tumor growth in PDCX slices, measured as the percent increase in GFP signal over time relative to baseline (n=58 slices). **(H)** Quantification of cell movement, measured as the percent spatial rearrangement of the GFP signal between consecutive frames (n=56 slices). **(I)** Log-log plot of mean squared displacement (MSD), where α represents the slope of the curve. A slope of 1 indicates Brownian motion (dotted line) (n=56 slices).

### GlioTrace detects divergent glioblastoma invasion dynamics across patient-specific models

Using GlioTrace, we analyzed PDCX brain slices from six patient cases, drawn from our in-house HGCC Phenobank collection. Recent work has shown that these PDCXs exhibit a diversity of growth phenotypes, forming two main groups defined by (i) bulky tumors with perivascular growth, and (ii) diffusely growing tumors with extensive white matter involvement [32]. Additionally, the collection contains rarer phenotypes such as significant meningeal growth. Despite previous studies documenting these phenotypic variations, the dynamic mechanisms driving these differences remained unclear. To address this gap, we analyzed six PDCX cultures derived from GBMs with different transcriptional subtypes (proneural, mesenchymal, and classical). Four of these—U3013MG (78y female, proneural), U3028MG (72y female, proneural), U3054MG (60y female, mesenchymal), and U3220MG (69y female, mesenchymal)—exhibited prominent bulky and perivascular invasion patterns **(Figure 1C, E, F, Supplementary Fig. 2A).** The other two—U3180MG (80y male, classical) and U3179MG (62y male, classical)—demonstrated diffuse invasion phenotypes, with extensive white matter involvement **(Figure 1D, Supplementary Fig. S2B).** Notably, U3220MG, derived from a gliosarcoma tumor, displayed strong subventricular and meningeal growth, while U3054MG, derived from a recurrent glioblastoma, showed a pronounced tendency for perivascular invasion **(Figure 1E–F).**

Time-lapse sequences revealed distinct invasion dynamics across the models. Tumor cells in all four bulky/perivascular models (U3013MG, U3028MG, U3220MG, and U3054MG) formed advancing edges where cells migrated collectively along vascular routes, confirming active perivascular invasion with perivascular cells engaging in both individual motion and collective stream-like motion **(Supplementary video 1B, 1C, 1D, 1E).** In contrast, cells in diffusely growing tumors (U3180MG, U3179MG) exhibited sporadic interactions with blood vessels, with predominantly individual cell migration **(Supplementary video 1F-1G)**. Interestingly, U3179MG cells migrated as discrete entities, regardless of vessel contact **(Supplementary video 1G)**.

To explore the patient specific differences in a systematic manner, we applied GlioTrace to measure three metrics of growth and invasion. First, quantification of GFP signals across the entire slice confirmed net tumor growth in five of the six PDCX models during the 6-day observation period **(Figure 1G)**. Growth rates varied significantly between subtypes, with gliosarcoma-like U3220MG and recurrent U3054MG slices exhibiting the most rapid growth, while other glioblastoma models showed slower growth rates **(Figure 1G)**. Second, to dissect cellular movement, we calculated the adjusted mean absolute deviation (adMAD) as a population-level metric of spatial rearrangement, accounting for signal changes due to growth. The highest adMAD values were observed in the recurrent U3054MG and primary U3028MG models, both of which demonstrated accelerating rates of cellular movement. Other models exhibited more consistent and moderate spatial rearrangement **(Figure 1H).** Third, to extend these statistics, we measured the movement by tracking individual cells within the slice, revealing a similar ranking of the different PDCX models, based on their generalized diffusion coefficient *D* as calculated from a Mean Squared Deviation (MSD) graph. Conceptually, the intercept of an MSD curve captures information about the overall diffusion rate of cells, and the slope α captures the nature of the motion, where a slope of 1 indicates Brownian motion and >1 indicates superdiffuse/directed motion **(Figure 1H)**. In our model, cells in bulky tumor models exhibited anomalous diffusion with larger intercepts, indicative of rapid, directed migration along vascular routes. In contrast, cells in diffusely growing glioblastoma models showed lower MSD intercepts, suggesting slower but still directed movement **(Figure 1I).**

Taken together, PDCX brain slices analyzed with GlioTrace demonstrated patient-specific invasion phenotypes, maintained viability, and exhibited clear tumor expansion during six days of observation. Bulk-forming tumors showed robust perivascular invasion, with coordinated migration along blood vessels at the invasive edge. In contrast, diffusely growing tumors displayed more sporadic vessel interactions and predominantly individual cell migration.

### GlioTrace reveals patterns of phenotypic state switching and migration route change

We went on to investigate different classes of migratory cells detectable by GlioTrace at the tumor periphery. Through the analysis of multiple regions of interest (ROIs), we identified three distinct migratory phenotypes. First, *branching cells* exhibited long cytoplasmic protrusions extending at various angles from the cell body, with PDCX models U3013MG and U3028MG frequently demonstrating this phenotype **(Figure 2A, Supplementary video 2A).** Second, *translocating cells* appeared spindle-shaped with central nuclei and forward and backward protrusions, often engaging with vasculature during migration **(Figure 2B, Supplementary video 2B).** Third, *locomotive cells* displayed unidirectional migration with elongated leading-edge protrusions and nuclei positioned at the rear. This phenotype was prominent in U3180MG, where cells often migrated along vascular routes **(Figure 2C, Supplementary video 2C).** Additional morphologies, such as round cells and atypical cell types, were also observed but occurred less frequently. These three phenotypes (branching, translocating, and locomotive) correspond relatively well to cell classes recently detected by 2-photon and 3-photon intravital microscopy setups [33, 34], and are here reported for the first time in a scalable slice culture system. In addition to these phenotypes, we noted cells with a round morphology, as well as atypical cells that did not confer to any of the above-mentioned morphologies.

**Figure 2.**
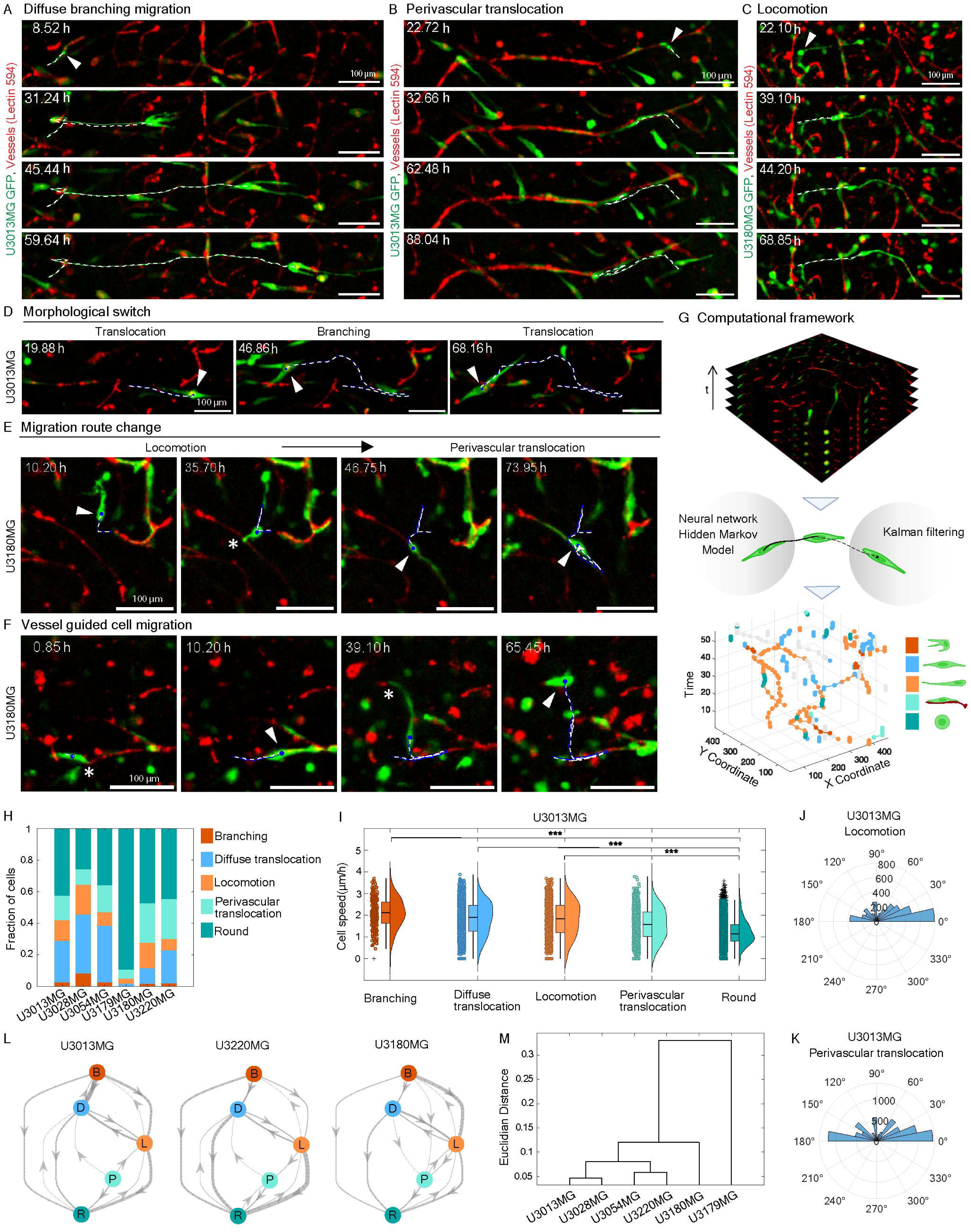
Migratory and invasive mechanisms of glioblastoma cells in PDCX slice culture. **(A, B)** Time-lapse confocal images of tumor cells displaying branching morphology and perivascular translocation. **(C)** Tumor cells exhibiting locomotion-driven migration. **(D)** Example of a glioblastoma cell undergoing morphological switching. **(E)** Tumor cell changing migration routes. **(F)** Example of vessel-guided migration. Arrowheads in A–E indicate tracked cells; asterisks in E, F indicate vessel contact. Scale bar: 100 μm. **(G)** GlioTrace computational framework extracting cell trajectories and morphological states. **(H)** Distribution of invasion phenotypes across PDCX models (n=56 slices). **(I)** Cell speed of different morphological classes in the U3013MG model (log-transformed values, n=21 slices, 18,716 cells). **(J, K)** Turning angle distributions of locomoting and perivascularly translocating cells in U3013MG (n=21 slices). **(L)** Hierarchical representation of state transitions, where thicker edges indicate more frequent transitions (n=41 slices). **(M)** Clustering of PDCX models based on state transition matrices.

Beyond these core phenotypes, we observed several intriguing dynamics. Tumor cells demonstrated phenotypic switching, transitioning between branching and translocating states, and reverting over time **(Figure 2D, Supplementary video 2D)**. Additionally, cells altered their migration routes, shifting between diffuse locomotion and perivascular migration **(Figure 2E, Supplementary video 2E)**. Notably, in U3180MG, diffusely invading cells alternated between vessel-associated and non-vessel-associated migration, indicating a highly adaptive invasion strategy **(Figure 2F, Supplementary video 2F)**. It thus appears that glioblastoma cells have an inherit ability to switch between modes of migration.

To analyze these dynamics systematically, we integrated a convolutional neural network (CNN) into the GlioTrace framework for phenotype classification and temporal tracking **(Figure 2G)**. The CNN, trained on 8,000 manually annotated cell images, achieved an 87% classification accuracy **(Figure S2E)** and categorized cells into distinct invasion phenotypes while identifying vessel associations. Analysis of over 3,000 ROIs confirmed that locomotive cells dominated U3180MG slices, while branching cells were most prevalent in U3028MG **(Figure 2H)**. Quantitative measures revealed that branching cells exhibited the fastest migration rates, while locomotive and translocating cells demonstrated the most directional movement, as indicated by turning angle distributions **(Figures 2I–K, Supplementary figure 2C).**

Using the time-sequenced phenotypic data of thousands of cells, we constructed Hidden Markov Models (HMMs) to infer phenotypic transitions and hierarchical relationships. Conceptually, the HMM models the morphological switching as a change in the hidden state of the HMM, and uses the emission probabilities to model the uncertainty of the CNN, thereby estimating a network of switching. The HMM revealed that branching cells served as a central transition state, while round cells often represented terminal or “sink” states **(Figure 2L, Supplementary figure 2D).** Clustering PDCX models based on their HMM parameters resulted in two distinct subgroups—bulky perivascular and diffusely invading tumors— consistent with manual observations **(Figure 2M).**

These findings underscore GlioTrace’s ability to comprehensively profile glioblastoma invasion phenotypes in a scalable format, characterize phenotypic transitions, and uncover adaptive migration strategies. When combined with an HMM model, this approach provides insights into the cellular hierarchies and dynamic behaviors that drive glioblastoma progression.

### Patient-specific vascular disorganization and activation of brain resident innate immune cells

Next, we investigated the power of GlioTrace in terms of observing functional interactions between glioblastoma cells and the surrounding brain. Glioblastoma cells are known to interact with the brain micro-vasculature, employing blood vessels as a scaffold for migration via the perivascular spaces [31, 35, 36].

A subset of our PDCX models exhibited extensive disorganization of the vasculature at the tumor front. In the U3013MG and U3028MG tumor slices, TL-DyLight 594-positive vascular structures were disrupted and retracted toward the tumor cell mass. Of note, vascular structures were disrupted even before physical contact between tumor cells and endothelial cells **(Figure 3A, Supplementary figure 3A, 3F, Supplementary video 3A & 3B),** suggesting mediation via a soluble factor. Similarly, U3220MG and U3054MG formed connections with vessels while disintegrating endothelial organization **(Supplementary figure 3C-D, H-I, Supplementary video 3C & 3D).** In contrast, while the diffusely growing U3180MG and U3179MG tumor slices also exhibited perivascular connections of tumor cells, this occurred without disruption of the vascular network and with the majority of tumor cells migrating independently regardless of vessel contact **(Figure 3B; Supplementary figure 3B, E, G, Supplementary video 3E & 3F).** To quantify these observations, we measured the extent of vessel disruption occurring around tumors by analyzing the length of vessels across the time-lapse, using a neural network for vessel segmentation. When comparing the length of vasculature at the tumor border to vasculature within control (tumor free brain) regions, we saw significant effects on vascular disorganization in all bulk-forming tumors and in one of the diffusely growing ones **(Figure 3C & 3D)**.

**Figure 3.**
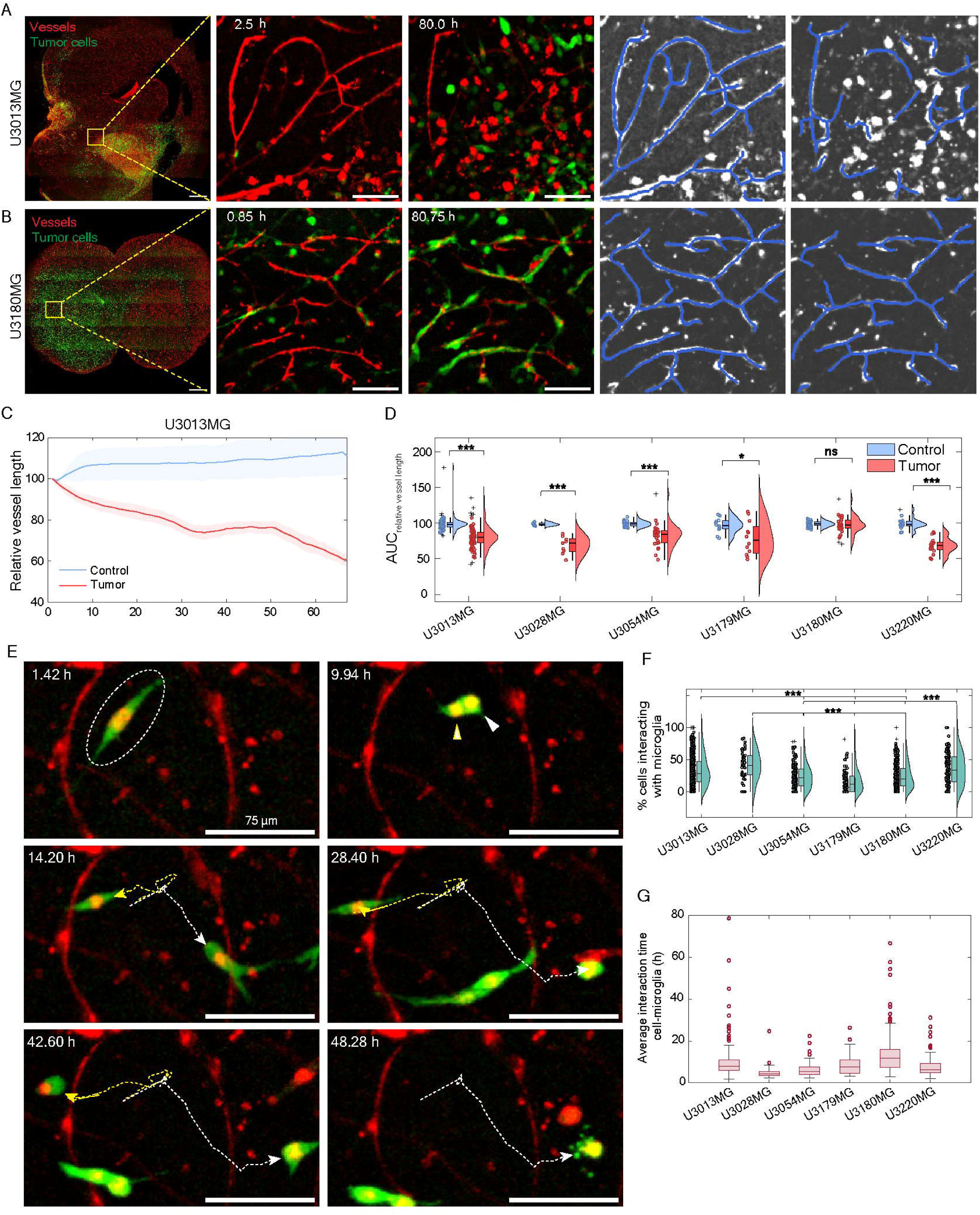
Vascular remodeling and immune interactions during glioblastoma invasion. **(A, B)** Whole-brain confocal images with zoomed-in time-lapse sequences (0 h and 80 h) showing vascular changes in U3013MG and U3180MG glioblastoma tumors. Blue overlay represents segmented vasculature. Scale bars: 1 mm (left), 100 μm (zoomed-in). **(C)** Comparative analysis of relative vessel length at the tumor front versus normal brain tissue in U3013MG slices (n=26 slices). **(D)** Violin plot showing area under the curve (AUC) of relative vessel length at the tumor border compared to control regions across PDCX models (n=61 slices). **(E)** Time-lapse images showing microglia-tumor cell interactions, with yellow and white arrowheads indicating synchronized glioblastoma-microglia division. Scale bar: 100 μm. Red: TL-DyLight 594; Green: GFP. **(F)** Percentage of tumor cells interacting with microglia/macrophages (n=56 slices). **(G)** Average duration of tumor cell-microglia/macrophage interactions (n=56 slices).

In addition to vascular structures, GlioTrace captures a population of relatively round and motile TL-DyLight 594-positive microglia/macrophage cells at the tumor core and periphery **(Figure 3E).** The TL-DyLight 594 signal correlates well with the microglia marker Iba1 protein staining at the tumor periphery, which, together with the fact that TL-DyLight 594 stain microglia [37], identifies this population as activated microglia **(Supplementary Figure 4A-C)**. GlioTrace, therefore, provides a means of probing microglia dynamics at the glioblastoma border. Intriguingly, we found examples of microglia seemingly ‘chasing’ tumor cells at the tumor periphery, ending with the death of the tumor cell **(Figure 3E; Supplementary video 3G).** Using a neural network trained to detect tumor cells co-localizing with microglia, we calculated the level of microglial engagement, defined as the percentage of tumor cells at the tumor periphery that co-localized with microglia, as well as the duration of colocalization. Broadly, the PDCX models with an ability to disorganize blood vessels showed higher levels of microglial/macrophage engagement **(Figure 3F)**. However, the relatively slow-moving U3180MG had a longer duration of microglial/macrophage engagement compared to the faster-moving cell lines **(Figure 3G)**.

These results highlight the utility of GlioTrace in observing the intricate spatial and temporal relationships between glioblastoma cells, vasculature, and immune populations, providing a robust platform to elucidate dynamic cell-cell interactions at the tumor’s edge region.

### Targeted drugs modulate invasion and phenotypic switching

Our observations that GlioTrace detects patient-specific trends in movement, phenotypic switching, vascular disorganization, and microglial interactions suggested a potential to explore how drugs affect these complex dynamics. To explore this capability, we focused on two drugs that were previously reported by us and demonstrated inhibition of glioblastoma cell migration and invasion in vitro: the Src/Bcr-Abl inhibitor dasatinib and the calcium channel blocker thapsigargin [38].

Using matched treated and untreated slices of U3013MG, we observed that both drugs elicited dose-dependent reductions in growth and invasion, evident through manual evaluation **(Figure 4A; Supplementary figure 5A–E; Supplementary videos 4A, 4B, 4C, 4D, 4E)**. Specifically, the average cell speed (µm/hour) decreased by more than 50% in all models except U3179MG **(Figure 4B)**. GlioTrace’s quantitative metrics—growth, movement, and diffusion—enabled a precise analysis of these effects using a linear mixed-effects model, revealing statistically significant dose-dependent treatment effects **(Figure 4C, D, Supplementary figure 5F)**. Notably, all models exhibiting reductions in speed also showed an increased proportion of round cells, indicative of a shift towards less invasive morphologies **(Figure 4E)**.

**Figure 4.**
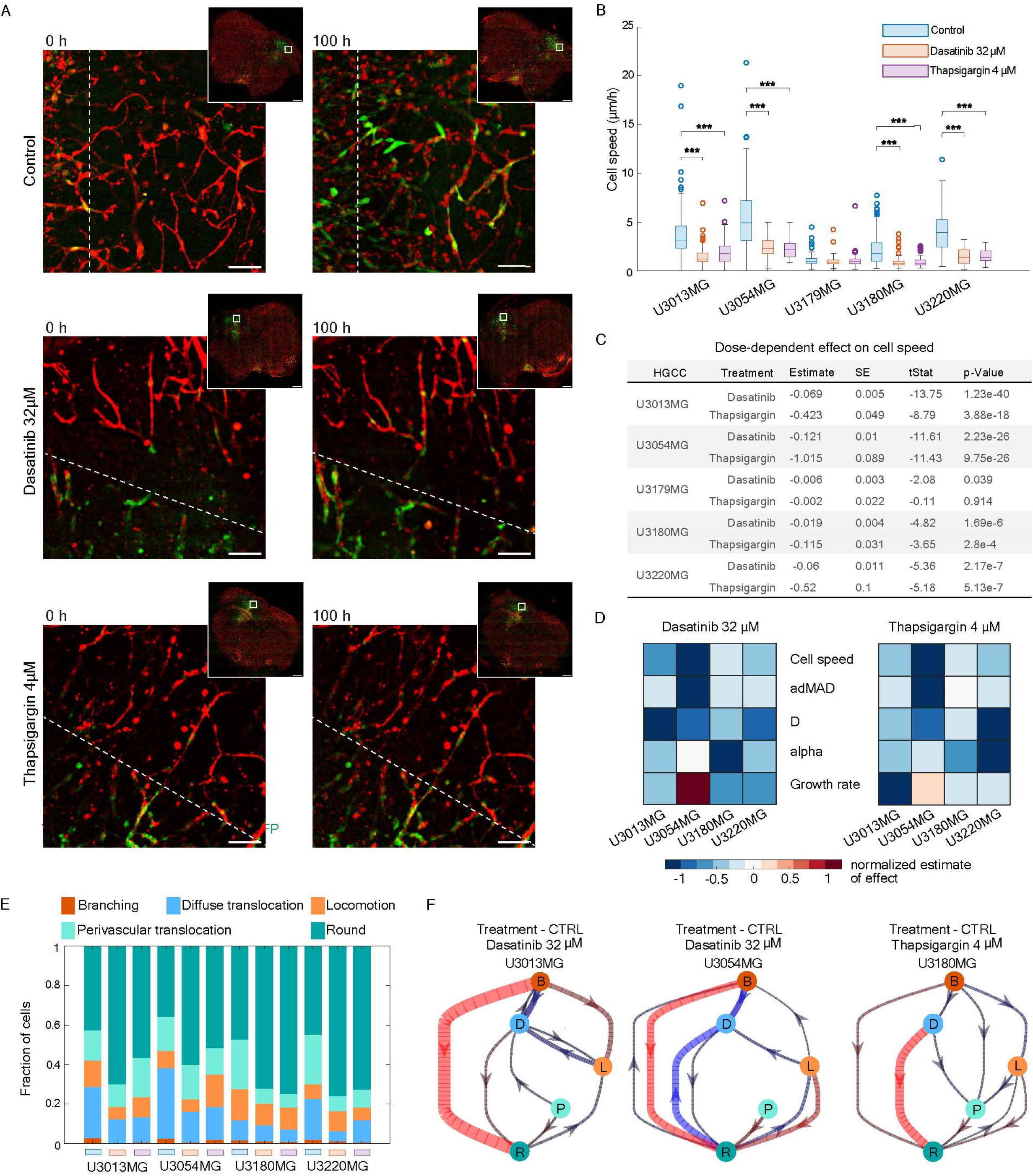
Drug effects on glioblastoma migration and invasion in PDCX slice cultures. **(A)** Representative confocal images of U3013MG slices at the first and last time points, untreated or treated with dasatinib (32 μM) or thapsigargin (4 μM). Tumor cells (green), vasculature (red). Dotted line indicates the initial tumor border. Scale bar: 100 μm. **(B)** Boxplot of cell speed across treatment conditions. (n=162 slices) **(C)** Summary of dose-dependent effects of dasatinib and thapsigargin on cell speed across PDCX models, estimated using a linear mixed-effects model (LME). (n=162 slices) **(D)** Heatmaps showing drug effects on five measures of movement and growth across PDCX models. Values are normalized LME estimates. **(E)** Morphological state distributions across PDCX models, comparing drug-treated and control conditions. (n=162 slices) **(F)** Differential state transition hierarchies for three PDCX models. Red and blue edges indicate increased or decreased transition rates, respectively, relative to control. Edge thickness reflects transition frequency (n=73 (U3013MG), 22 (U3054MG), and 36 (U3180MG) slices).

Branching cells, often associated with rapid invasion **(Figure 2I)**, were particularly sensitive to drug treatments. Dasatinib and thapsigargin both completely eliminated branching cells in U3013MG, whereas U3220MG retained branching cells after dasatinib treatment but lost them following exposure to thapsigargin **(Figure 4E)**. To capture the underlying dynamics, we used our Hidden Markov Model (HMM) framework to work out differential morphological state transitions upon treatment. Across all models, transitions to the round phenotype—interpreted as indicative of cell death—were predominant. However, drug-specific effects were also evident. For example, dasatinib-treated U3054MG cells showed a reduction in locomoting-to-diffuse translocation transitions, highlighting the drug’s selective inhibition of certain invasive phenotypes **(Figure 4F, Supplementary figure 4G)**.

Together, these results show that known inhibitors of invasion produce statistically strong effects, and that the ability of GlioTrace to extract multiple metrics suggests that it may be used to group pharmacological agents into multiple classes based on their effect profile. We also propose that GlioTrace be used as a tool to quantify the rewiring of morphological state transitions and selective killing of specific migratory phenotypes, following both chemical and genetic perturbations.

### Mapping spatiotemporal trends in growth, movement, and drug response

Continuous monitoring of the whole specimen provides a novel means to map spatial and temporal variations in tumor behavior, enabling insights into both growth dynamics and drug-induced changes. Using 3D image stacks, we applied a smoothing kernel estimator to analyze tumor growth and adMAD, our measure of pixel turnover reflecting cellular movement, in untreated and drug-exposed slices.

As expected, tumor growth was centered in the tumor core, with adMAD indicating significant cellular motion concentrated at the periphery **(Figure 5A, B).** Post-imaging Ki67 protein staining, which marks proliferating cells, revealed that the motile cells at the periphery were largely non-proliferative, displaying morphologies consistent with invasive phenotypes **(Figure 5C, E)**. Exploring the effects of dasatinib, we observed a marked reduction in Ki67-positive cells within the tumor core, indicating a significant anti-proliferative effect in U3013MG slices **(Figure 5C, Supplementary figure 6A).** The reduction in core growth was consistent with spatial growth estimates, which showed the greatest decline in proliferative activity at the tumor center. However, dasatinib-treated U3054MG slices maintained peripheral motility at reduced levels, suggesting a differential impact on proliferative and invasive phenotypes. **(Figure 5D, F)**. In contrast, thapsigargin treatment failed to inhibit core proliferation in both models, as evidenced by persistent Ki67 positive cells **(Figure 5F, Supplementary figure 6A-B).**

**Figure 5.**
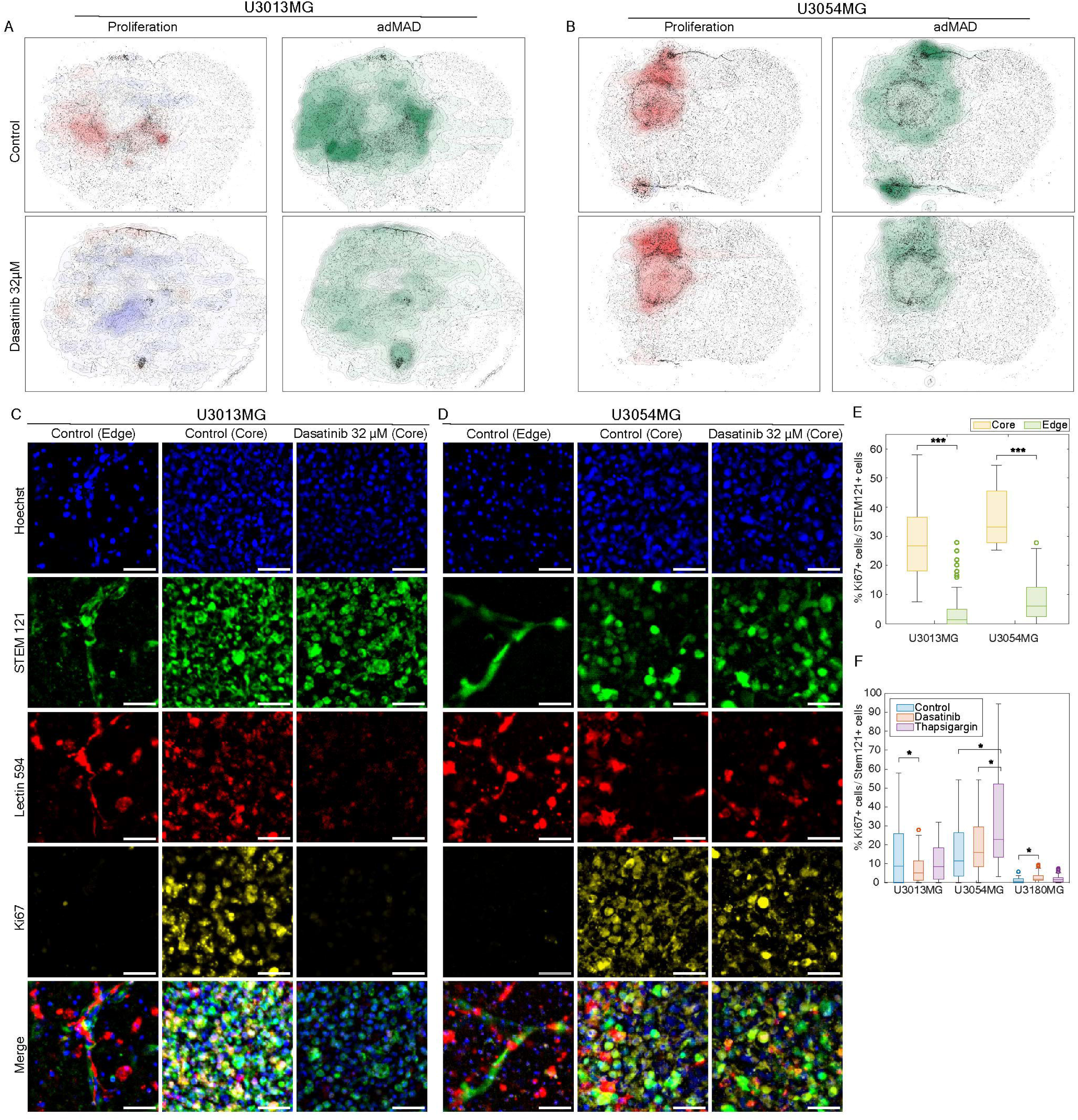
Spatial analysis and post-timelapse immunostaining reveal differential drug effects in tumor core and invasive front. **(A, B)** Spatial maps of tumor growth and migration in whole brain slices, illustrating differential drug responses in tumor core and invasive front. Blue: reduced growth, Red: increased growth, Green: migration extent. **(C, D)** Confocal images of the tumor invasive front in U3013MG and U3054MG slices stained for Ki67 after time-lapse imaging. **(E)** Percentage of Ki67+ proliferating cells normalized to STEM121+ tumor cells, comparing core and edge regions in U3013MG and U3054MG (n=36). **(F)** Ki67+ tumor cells in dasatinib- and thapsigargin-treated slices, normalized to control (n=36).

Taken together, GlioTrace can resolve the zonation of a growing tumor and estimate effects on growth consistent with a standard histological marker. The spatial separation of proliferative and invasive behaviors underscores the heterogeneity of glioblastoma dynamics.

## Discussion

Glioblastoma invasion is emerging as a complex process where heterogeneous cell populations transition between different invasive phenotypes. Exploring these phenomena at scale requires a high-resolution observational platform and the capability to interpret individual cell dynamics across patient-specific (diverse) tumor microenvironments. Our methodological advancement meets the need for a system that can resolve complex spatiotemporal phenotypes in grafted tumors in a scalable format with moderate cost. This method can be implemented on standard automated confocal microscopy equipment and with modest computing resources. Potential applications include exploring modulation of the tumor-microenvironment interactions, testing of new anti-migratory treatments, patient stratification, and comparing the dynamics of different zones of the tumor.

Our comparative exploration of a set of PDCX models from the HGCC collection underscores the complexity of brain tumor invasion. In addition to detecting invasive phenotypes observed in more complex animal experiment setups [33, 34, 39], GlioTrace makes a number of intriguing findings. First, we confirm that GBM cells exhibit dynamic transitions between different migration routes, including shifts between diffuse locomotion and perivascular translocation. These transitions appear to be influenced by both intrinsic cellular states and interactions with the microenvironment, highlighting a level of invasion plasticity that warrants further investigation. Second, we distinguish between translocating and locomotive invasion based on distinct nuclear positioning and cellular morphology, suggesting that these modes may be regulated by different molecular mechanisms. Moreover, we identify a highly motile branching cell type that frequently transitions between invasion phenotypes, potentially representing a key top state in GBM invasion hierarchies. Finally, our observation that vascular disorganization and microglial/macrophage engagement are more pronounced in proneural and mesenchymal tumors raises the possibility that microenvironmental interactions contribute to subtype-specific invasion strategies. Further studies are needed to determine whether targeting these interactions could provide new therapeutic avenues for controlling GBM invasion. Linking the dynamics detected by GlioTrace with underlying molecular processes is a key priority for future research; this will require refined histological analysis of brain slices following GlioTrace imaging, spatial molecular profiling strategies, or conditional reporters. While the number of patient cases explored in this study is limited to six, we point to the fact that the two diffusely invading cultures (U3180MG, U3179MG) are of a classical subtype, with strong expression of astrocyte markers [32]. Moreover, we recently found that the translocating cell portion of U3013MG are mesenchymal in nature, depending on ANXA1 for their distribution along blood vessels [32].

GlioTrace complements existing assays through its scalability and observational power. Working across a broad field-of-view, it detects multiple types of invasive cells, scores cell-cell interactions, and quantifies patient differences as well as drug effects. Compared to traditional slice culture [26, 31, 40], GlioTrace provides more comprehensive spatiotemporal observation and analytical depth. Compared to intravital microscopy using 2- or 3-photon setups [34], it scales well and is fraught with fewer practical and ethical concerns. The benefit of the scalability becomes clear when considering that we detect clear patient differences in both growth, migratory cell morphologies, invasion routes, cell-cell-interactions, and drug responses. Compared to increasingly popular organoid model systems, the presence of a reference mouse brain anatomy, vascular and glial components will be an advantage for specific applications.

GlioTrace must be used with an awareness of the inherent limitations of brain slice assays and restricting the analysis to two optical channels. Being on an air-liquid interface, the brain slice assay will not emulate the Blood-brain barrier (BBB), and the system will also carry inherent limitations of being a mouse-human hybrid without other myeloid cells and lymphoid cells present. A second technological limitation is the current approach of projecting multiple Z stack imaging layers into a max intensity 2D projection image for analysis at each time point. In some cases, especially in the tumor’s core region, this will limit our ability to segment individual cells. However, at the edge region, the method provides good separation, both between individual tumor cells in the green channel, and between microglia/macrophage vs vascular structures in the red channel. The neural network applied here gives excellent separation of classes (87% for 6 classes, i.e. a null expectation of 17%) comparable to similar classification problems [41] and any remaining uncertainty is accounted for by the HMM. However, we do not exclude that some users may need to retrain or fine-tune the network depending on their exact imaging setup or other cells of interest.

Moving forward, we see a number of promising extensions to our approach. First, we are seeking to extend the labeling strategy to comprise multiple cell types and markers, which is currently easy to implement on many microscopes, coupled with the extended single-cell tracking of such cell types. This would, for instance, enhance the system’s ability to score specific migratory phenotypes such as white matter penetrating cells, or glioneuronal interactions. A second important extension is to expand cell tracking into a full 3D setting. Last, as vascularized CNS organoids are increasingly robust, we foresee an adaptation of GlioTrace into a human-only model setting, with potential extensions to the evaluation of drug responses in clinical samples.

Practically, GlioTrace can now be distributed as an open-source analysis pipeline to be used for any set of ex vivo immunofluorescence images acquired through standard live confocal or brightfield microscopy with 10X magnification or greater (Code and data availability). GlioTrace could be adopted or trained further to generate all types of analysis performed in this study for multiple cell types with separate fluorescence tags. Taken together, we expect GlioTrace to enhance and expedite the discovery of anti-migratory treatments for glioblastoma, with possible applications in other tumors.

## Methods

### Models used in experiments and subject details

The mouse experiments described herein were carried out in accordance with an ethical permit granted by the Uppsala Animal Research Ethical Board (Ethical permit number: 5.8.18-06726/2020 and 5.8.18-00216/2023). Female mice from 5 weeks of age (Hsd: Athymic Nude-Foxn1^nu^, Envigo) were housed in ventilated cages, each accommodating up to five mice. These cages were equipped with suitable paper-based enrichment and bedding material, while the mice had unrestricted access to food and water. Environmental conditions were controlled to adhere to a 12-hour light/12-hour dark cycle.

### GFP-tagged human glioblastoma patient-derived primary cancer cell cultures

Primary cultures of patient-derived tumor cell lines as outlined in supplementary data, were procured from the Human Glioblastoma Cell Culture (HGCC) biobank [42, 43]. The collection of HGCC samples was authorized by the Uppsala Regional Ethical Review Board (2007/353), and explicit informed consent was obtained from all participants. These cells were propagated in laminin-coated T25 flasks (Corning #353808) utilizing a serum-free neural stem cell culture (NSC^+/+^) medium, comprising 50:50 nutrient blend of Neurobasal and DMEM/F12 media (Gibco, #21103-049, #31331-028), supplemented with B27 and N2 additives (ThermoFisher Scientific, #12587-001, #17502-001), human recombinant epidermal growth factor (EGF) and fibroblast growth factor (FGF) at concentrations of 20 ng/ml each (Peprotech), along with 1% penicillin/streptomycin. All utilized cell cultures were previously transduced with Green Fluorescent Protein (GFP) and luciferase (pBMN(CMV-copGFP-Luc2-Puro), Addgene plasmid #80389) [44]. Furthermore, all cell lines underwent scrutiny for mycoplasma contamination using the Mycoalert kit (Lonza, #LT07-418) and were affirmed to be devoid of such contamination.

### Xenotransplantation of patient-derived glioblastoma cells in mice

Each GFP-labeled HGCC line was prepared at a density of 1 million cells per 20 µl in phosphate-buffered saline. Mice aged 6 weeks were anesthetized using deep isoflurane inhalation within an induction chamber and subsequently secured onto a stereotactic frame under anesthesia. Subsequently, 100,000 tumor cells were stereotactically injected into the striatum of the mouse brain at the following coordinates relative to the bregma: AP +0, ML 1.5(R), DV -3.0. Mice were closely monitored until fully awake, and post-injection weights were recorded to monitor weight changes throughout the experiment.

### Glioblastoma PDCX tumor growth monitoring through bioluminescence

Following 6 weeks post-implantation of glioblastoma into the striatum using stereotactic techniques, mice received an intraperitoneal injection of 100µl of luciferin solution (15µg/ml), followed by *in vivo* IVIS spectrum imaging (Perkin Elmer, Waltham MA) every 2 weeks to quantify luciferase signal levels indicative of optimal tumor growth.

### Vibratome sectioning and PDCX brain slice culture

Intermittently, when luciferase levels indicated optimal tumor growth (or size), mice were euthanized individually, and their brains were aseptically extracted and submerged in 50 ml of cold HBSS medium (Gibco™, #24020117) to facilitate preparation for vibratome sectioning. The brain was then encased in a low-melting agarose-HBSS medium cube. Organotypic brain slice cultures, each 300 µm in thickness, were generated through coronal sectioning of the brain utilizing a Leica vibratome (Leica VT 1200 S) instrument, with real-time sectioning speed adjusted between 20 – 200 µm/second to accommodate variations in brain texture, particularly in areas affected by tumor growth. Subsequently, brain slices were immersed in ice-cold brain slice culture medium (NSC +/+, 2.5 mM HEPES, 10 mM glucose) within a 6 cm culture dish. These slices were then transferred onto transwell membranes in a 12-well plate (Corning, #3460), with each well containing 700 µl of brain slice culture medium. Excess medium surrounding the brain slices inside the transwell was carefully removed to maintain the air-liquid interface necessary for optimal brain slice culture conditions. The brain slice cultures were then placed in a 5% CO2 incubator at 37°C for 24 hours to recover and acclimate from any physical trauma incurred during the slicing process.

### Time-lapse whole brain slice imaging and post-acquisition processing

Time-lapse Z-stack images of brain slices were captured using the ImageXpress Micro Confocal High-Content Imaging System (IXMCS) (Molecular Devices), with imaging parameters set to a Z-step size of 10 µm, 9×9 tiles, a 10X objective with calibration of 1.36 µm/pixel. These parameters were adjusted to ensure optimized imaging frequency, with intervals of no more than 2 hours between frames. The brain slice culture medium was replenished every 48 hours, and imaging sessions continued for up to 5 days. Subsequently, all brain slice culture images from each experiment underwent processing and montage creation using the montage and overlay journal in MetaXpress 6.5 software (Molecular Devices). Following image tiling, sequences were stack registered and aligned for frame-to-frame coherence using custom-written MATLAB (The MathWorks) scripts, facilitating statistical analysis of cell dynamics and tumor growth.

### Tomato lectin-DyLight 594 conjugate bioavailability assay

The bioavailability of the compounds in slice cultures was evaluated by a tissue panetrant conjugate TL-DyLight 594 (ThermoFisher Scientific, #L32471). Brain slices were incubated with brain slice culture medium containing TL-DyLight 594 at a concentration of 2 µg/ml. Subsequent to the addition of TL-DyLight 594, time-lapse images were captured at intervals of 1.4 hours over a 24-hour period.

### Evaluation of slice viability

To assess PDCX slice viability, brain slices underwent culturing for 0, 1, 3, and 6 days. Following each incubation period, slices were incubated with 5 µM Hoechst 33342 (ThermoFisher Scientific, #62249) in the brain slice culture medium, a cell-membrane permeable dye that labels all cells in the sample, and 2 µM EthD-1 (Sigma-Aldrich, #46043-1MG-F), a live cell-impermeable dye that selectively labels dead cells, was applied for 45 minutes at room temperature (RT) by gentle shaking. After incubation, brain slices were washed with fresh brain slice culture medium and imaged using the IXMCS. The slice viability was quantified as the percentage of EthD-1 positive cells out of the total number of Hoechst cells for 2D projection images saved as maximum projection of 15 z-stacks taken at 10 µm interval.

### Ex vivo drug treatments

For the in vitro drug treatment studies, Four HGCC cell lines (Supplementary data) seeded at a density of 2500 cells per well in a 96 well Primaria plate (Corning, #353872) coated with laminin and grown in a 5% CO2 incubator at 37 °C. The next day, dasatinib (Selleck chemicals, #S1021) and thapsigargin (Sigma-Aldrich, #T9033) treatments were added in the concentration range of 0.005, 0.05, 0.5, 1, 2, 4, 8, 16, 32, and 50 µM. Plates were then immediately transferred into an Incucyte S3 (Sartorius) for live cell image acquisition every 30 min for 72 h.

For the PDCX brain slice culture treatment studies, slices were treated with dasatinib and thapsigargin at a concentration of 32 µM and 4 µM, respectively, prepared in brain slice culture medium. For a dasatinib dose escalation treatment PDCX brain slices were treated with 0.05, 1, 4, 8, 16 and 32 µM dasatinib. For the thapsigargin dose escalation treatment, PDCX brain slices were treated with 0.05, 1, 2, and 4 µM thapsigargin. Soon after adding the drug treatment, slices were transferred to the IXMCS system for time-lapse image acquisition.

### Slice immunofluorescence staining

Upon completion of time-lapse image acquisition, brain slices were fixed with 1-2 ml of freshly prepared 4% paraformaldehyde (PFA) for 2 h at room temperature (RT) and subsequently transferred into a 24-well plate. Following a triple wash with phosphate-buffered saline (PBS), the brain slices were subjected to incubation in a blocking solution (comprising 3% normal goat serum and 0.3% Triton X-100 in PBS) for 2 hours at RT. Thereafter, slices were exposed to primary antibodies, including mouse anti-STEM121 (diluted 1:500; Takara Bio, #Y40410) and rabbit anti-Ki67 (diluted 1:300; Abcam, #ab15580), prepared in the blocking solution. Following a 2-day incubation period, the slices underwent triple PBS washes and subsequent incubation with appropriate Alexa Fluor-conjugated antibodies for 4-5 hours, followed by another triple PBS wash. Hoechst staining was then performed for 30 minutes.

### Image registration and ROI selection

The input data from the IMX system were provided as 9000×9000 whole slide scans for a sequence of time points 1, 2,…, T. The image sequences were stabilized using rigid (translation + rotation) affine transformations based on automatically extracted landmarks [45]. Regions of interest (ROIs) were extracted from the slice to create stacks of size W x W x T where W is the size of the region. ROI selection was followed by stabilization using an efficient rigid stack registration algorithm based on a 2-dimensional Fast Fourier transform [46].

### Detecting tumor cell proliferation and movement

1. ROI-level summary measures. As a first layer of quantification, we computed ROI-level measures of growth and movement. To measure growth, a single region was chosen for each slice that would encompass the full tumor region (unless multiple tumor sites were present in the slice, in which case multiple ROIs were selected to accurately capture the full tumor). For each stack or collection of stacks representing the whole tumor, the total pixel intensity of the green channel in each frame is normalized by region size and divided by the total pixel intensity of the first frame in the stack to generate a sequence of values of length T, corresponding to the percentual increase in green channel signal over time. To detect movement, multiple smaller regions from each slice were chosen. Regions were primarily selected to be suitable for single-cell tracking, avoiding tumor bulk or empty regions. For each stack from the peripheral tumor regions in the slice a time-resolved adjusted mean of absolute differences (adMAD) was calculated according to Equation 1 where g is the green channel in stack G of size W x W x T. The absolute difference in mean pixel intensity between two frames was normalized by the mean pixel intensity of the two frames together to adjust for an increasing number of cells in the frame over time. The resulting measure describes the rearrangement of pixels between frames over time.

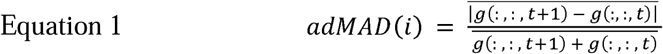

2. Cell tracking. To track single cells, we continued working with the peripheral ROIs suitable for this goal. Individual cell center detection was combined with a Kalman filter-based framework for multi-object tracking. Cells were detected with means of common image analysis approaches (Image Processing Toolbox, Computer Vision Toolbox, Version 23.2, Release 2023b, The MathWorks, Inc., Natick, Massachusetts, United States) intended to find intensity peaks in an image that could correspond to cell centers (Algorithm 1). Generated peaks were filtered through a number of steps to ensure that peaks (coordinates) were located to cell bodies (not cell protrusions), that they corresponded to single cells and that one cell had only one coordinate. The pruning of candidate points was followed by feature extraction; for each coordinate an image of window size M x M x C was extracted from the original image stack frame, where M is the size in pixels and C is the number of channels.

Cell coordinates and corresponding images are passed on to a multi-object tracking framework (Algorithm 2). The tracking framework is based on the Kalmar filter algorithm which connects spatial coordinates across time to create object trajectories. It relies on assumptions of an object’s velocity and acceleration, the dynamic model. Based on the dynamic model, coordinates are extrapolated into the near future and become predictions. The predictions are used to match coordinates between two subsequent frames by solving a linear assignment problem. The resulting tracks consist of coordinates x(t), y(t).

3. Summary statistics. From the tracks, we computed a number of summary statistics. These were the average speed (microns per hour) averaged across the whole track and the turning angle distribution obtained by standard trigonometry functions. Furthermore, the mean squared displacement (MSD) was calculated from the cell trajectory coordinates. For each stack, every cell trajectory is handled separately to calculate the squared displacement for each coordinate from the starting point of the trajectory. The tracks are interpolated to the smallest Δt and the final MSD is calculated as the ensemble mean across all trajectories from all stacks cell line-wise. Tracks are handled in such a way that they all start at t_0_, so that the time axis of the MSD curve corresponds to the time elapsed from the start of a track and not to the time elapsed in image acquisition.

### Detecting tumor cell morphologies and phenotypic switching

1. Morphological classifier. The images extracted around cell coordinates in the cell detection step were classified as belonging to one of six classes using a convolutional neural network (CNN) (Deep Learning Toolbox, Version 23.2, Release 2023b, The MathWorks, Inc., Natick, Massachusetts, United States). The network was trained on 7800 manually curated images of the 6 morphological classes. The initial dataset had some class imbalance, which was handled by doing a stratified split into training, validation and test sets, as well as oversampling classes in the training set.

Mean subtraction centering was performed by subtracting the mean training image from all images in both training, validation and test set channel-wise. Mean subtraction centering with the same mean image was also performed channel-wise when images were classified inside the framework. Augmentation of training and validation images was achieved with the MATLAB function augmentedImageDatastore and included reflection, rotation and scaling.

2. Hidden Markov Model (HMM) and cell state calling. By applying the above classifier to each cell along its track, we obtained a sequence of class assignments C(t), where t=1, 2, …, is the time step along the track. Our HMM views the class sequence C as produced by a hidden Markov model with hidden state transition matrix A and an emission probability matrix B. Specifically, we view the hidden state as the “true” biological state of the cell, while the class assignment becomes the observed (assigned) state. The full model is specified by

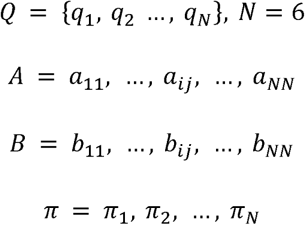

where Q is the possible true states of the cell, A is a matrix of hidden state transition probabilities, B is a matrix of emission probabilities, and π is the starting state probability distribution. Applied to our problem, the entries of A={a_ij_} will give the probability of proceeding from state i at time t to state j at time t+1. The entries of B={a_ij_} in turn give the probability for a cell that is truly in (hidden) state i to be assigned the class j by the neural network, i.e. the classifier confusion matrix standardized to column sum 1. For a perfect classifier, B would be the identity matrix. Here, we estimate B as the confusion matrix when evaluating the CNN classifier on a test set **(Supplementary figure S2E)**.

Given a set of observed state sequences C1(t) C2(t), …, CM(t) for M tracks, our learning task is to estimate the unknown state transition matrix A and initial state distribution π. We do so by a modified version of Baum Welch’s algorithm, keeping B constant, and introducing a sparsity regularization for A. For robust results, we run the algorithm 1000 times, and use the average A across runs. Once fitted, the model can be used to de-noise the observed state sequences using Viterbi’s algorithm. For each track, the algorithm returns the hidden state sequence that yields the observed sequence most probable, resulting in a final state call for each cell at each time point. These, in turn were used for estimating the morphological cell state fractions throughout the paper.

### Detecting microenvironmental interactions

1. Blood vessel disorganization. In matched tumor-free and tumor-peripheral regions, we quantified the length of the vasculature in every frame of the image stacks. To the red (TL-DyLight 594) channel of these stacks, we applied semantic segmentation by a U-net trained on 400 manually labeled 250×250 images of vasculature. Five-fold cross-validation gave an f1-score of 0.74 ± 0.015, demonstrating that the network can learn well to segment vessels and generalize to unseen data. The binary segmentation mask created by the network was skeletonized using the medial axis transform and pixels were summed up to calculate the length of the vasculature. The time-resolved measure of vasculature length was normalized to mean of control curves and to t_0_. From the curves, we computed a vascular disorganization index as the area under the curve, normalized by the stack-length.
2. Classifying tumor-microenvironmental interactions. To identify tumor cells colocalizing with microglia or migrating along blood vessels in the images, we developed a classifier using the same workflow as described above and a curated dataset of 3000 manually labeled images. The network scored the images as belonging to one of the two classes, thresholding the score with class-specific thresholds. Hence, if the classifier is too unsure of either class, the cell is classified as ’non-associated’. The network achieved a performance of 92% accuracy on the test set **(Figure S3F)**. The class sequences obtained were used for calculating the percentage of cells interacting with microglia and the average duration of this interaction.

### Assessing dose-dependent effects on cell statistics and morphological transition rates

1. Detecting patient differences and treatment effects. The above framework can be thought of as a nested design that results in summary statistics (speed, MSD, or morphological class frequencies) at the level of a particular cell, region of interest, slice, mouse brain, or patient, with treatment as an additional experimental factor. To perform statistical testing, we therefore use linear mixed effects models (Statistics and Machine Learning Toolbox, Version 23.2, Release 2023b, The MathWorks, Inc., Natick, Massachusetts, United States). To detect the effects of drugs we used (in Wilkinson notation) Y ∼ treatment:dose + (1|batch), where each batch is a collection of slices cultured at the same time. We acknowledge that further study is needed to accommodate for non-linear responses to drugs and reserve such extensions to future work.
2. Differential transition matrices. As an additional explorative statistic, we calculated the effect of treatment on the hidden transition matrix A as simply the treated A matrix minus the negative control A matrix.

### Inferring spatial differences in drug response from whole-slice analysis

To identify spatial trends in the drug response, we applied growth- and movement metrics to the entire slice, applying it to both tumor peripheral and core regions. Downsampling of the stack w.r.t frame size was followed by Gaussian and Notch filtering. We estimated the growth across the slice as the median rate of log2 fold-change in GFP signal over time in the stack. Movement was estimated as the absolute difference between consecutive frames t+1 – t normalized by abs(t) and averaged across time to create a mean absolute difference in GFP signal over time above some threshold.

### Cell number quantification in IHC stainings

For IHC stainings, we applied Gaussian filtering followed by blob detection by convolution with a Logarithm of Gaussian-filter corresponding to average cell size. Potential cell coordinates found in the blob detection were pruned by using the filtered, binarized version of the staining as a mask.

### Interpolation of data

When dealing with data from image stacks acquired with different time intervals between frames (Δt), data points were interpolated to the smallest Δt with the MATLAB function interp1.

### Pseudocode

#### ALGORITHM 1: Cell Center Detection

1. For each frame f = 1, 2, 3, …
2. Perform blob detection by convoluting the green channel with a Laplacian of Gaussian filter to find local maxima
3. Perform morphological opening of a binary version of the convoluted image to create a mask representing cell bodies
4. Using MATLAB functions bwlabel and regionprops, threshold cell body objects to remove objects above some area threshold
5. Create a new mask by smoothing and binarizing the green channel with a Gaussian filter followed by imbinarize
6. For each of the labelled cell body objects
7. Find the local maxima present within the cell body object and select only those maxima which are also present in a binarized version of the convoluted image and the binarized green channel.
8. If the maximum intensity of the Gaussian filtered mask exceeds some threshold, exclude local maxima present within this mask.
9. If there are still several local maxima present, create a mean coordinate. If this mean coordinate is within the cell body and the area of the cell body is within some upper and lower boundary, keep this mean coordinate instead. Otherwise, keep the original local maxima.
10. End for loop
11. Use the pruned cell coordinates to extract images of size B x B with the extractFeatures function. Coordinates that are too far out along the image borders and cannot be used to extract images are discarded by only saving the coordinates returned by extractFeatures.
12. End for loop

#### ALGORITHM 2: Multi-object tracking with Kalman Filtering

1. Initialize track estimates x1,…,xk as the detections in frame 1 with velocity zero.
2. Let xf-1 be the extrapolated initial track estimates.
3. For each frame f=2, 3, …
4. Let q be the coordinates of the cells in the current frame
5. Pair coordinates and extrapolated predictions by using MATLAB function matchpairs where the cost matrix is D(xf-1, q)
6. Perform a state update of the paired tracks
7. Extrapolate the state
8. Extrapolate the coordinates of new tracks that didn’t get a corresponding old match
9. For old tracks that didn’t get a corresponding new match, add a penalty of +1 to a score counter.
10. Tracks with a penalty score over T are discarded.
11. End for loop

### Statistical Analysis

We performed at least two independent experiments with minimum two biological replicates. Pairwise comparisons were performed with one-way ANOVA and, when involving multiple pairwise comparisons, with Tukey’s HSD test, with a significance level of 0.05.

## Acknowledgements

We thank the patients and their families for the support and donation of materials to the HGCC biobank. We thank Lioudmila Elfineh for providing GFP labelled HGCC lines used in this work. We thank our colleagues Veronica Rendo, Bengt Westermark, Fredrik J Swartling and Karin Forsberg Nilsson at the Uppsala University program Neuro-Oncology and Neuro-Degeneration for their valuable comments on the first draft of the manuscript.

## Author contributions

Conceptualization by SN, CK, SK, HBM and MS. Formal analysis, investigation and validation by HBM and MS. Methodology by HBM, MS, CK and SN. Visualization by HBM, MS and SN. Animal handling by HBM and MB. Software by MS, SN and AM. Writing – original draft by HBM. Writing – review and editing by SN, HBM and MS. Funding acquisition and supervision by SN.

## Funding

This research work was supported by the Swedish Research Council (2021-03224), Swedish Cancer Society (23-0696-PT), and the Knut och Alice Wallenbergs Stiftelse (2022.0057)

## Data and code availability

The whole brain slice images from each of the HGCC lines could be made available upon request. The code base for the GlioTrace framework and additional image processing and analysis scripts is available as a GitHub repository (https://github.com/shipsauce/brainslice_manuscript_repo).

## Reporting summary

Further information on research design is available in the Nature Portfolio reporting Summary linked to this study.

## Declaration of interest

The authors declare no conflict of interests.

## Declaration of generative AI and AI-assisted technologies in the writing process

All text was reviewed and edited by the authors, and the authors take full responsibility for the scientific content written in this manuscript for the publication. The spelling and grammar of the manuscript were checked using Microsoft Word (Microsoft, Inc.), GPT4o (openAI, Inc.), and Grammarly (Grammarly, Inc.).

## Figure legends

**Supplementary figure 1. Validation of TL-DyLight 594 bioavailability to vessels and relative slice viability in longitudinal culture. (A)** Representative time-lapse confocal images showing real-time lectin-594 staining of vessels (red) in live U3013MG tumor (green) slice culture. **(B)** Percent pixel area of lectin-594 stained vessels in slice culture with or without tumor (n=4 slices). **(C)** Representative confocal images stained with dead cell membrane permeable dye EthD-1(red) and nuclear counterstain Hoechst (blue) at 0 day and after 6 day glioblastoma PDX brain slices in culture. Tumor cells are green. Scale bar: 1 mm (left column). Scale bar: 100 µm (zoomed) **(D)** Quantification of cell viability in the ex vivo U3013MG brain slice culture (n=12 slices).

**Supplementary figure 2. Demonstration of PDXs recapitulating human glioblastoma phenotype in slice culture.** Representative first and last time point confocal images showing **(A)** perivascular migration by proneural type (U3028MG, 72y female) glioblastoma tumor cells and **(B)** diffuse migration by classical type (U3179MG, 62y male) glioblastoma tumor cells in slice culture. **(C)** Turning angle distribution of migratory cells demonstrating branching, diffuse translocation, round and junk (cells with an unidentifiable morphology). **(D)** State switching hierarchies displaying the transition rates between different morphological states of U3179MG, U3028MG, and U3054MG tumor cells. Thicker line signifies a more frequent transition. (E) Confusion matrix of the neural network that classifies cells by morphology. B-Branching, DT-Diffuse Translocation, J-Junk, L-Locomotion, PT-Perivascular Translocation, R-Round.

**Supplementary figure 3. Patient specific tumor-vessels interaction in slice culture. (A - D)** Confocal whole-brain slice images at the starting point (left) with zoomed-in snippets at 0 h and end point of time lapse imaging highlight structural change in tumor associated vasculature in proneural (U3028MG), classical (U3179MG), and mesenchymal (U3220MG, U3054MG) glioblastoma tumors. Scale bars: 1 mm (left), 100 μm (zoomed). **(E-I)** Comparative measurement of relative vessel length at tumor progressing edge and normal brain area from same slices in U3180MG, U028MG, U3179MG, U3220MG, and U3054MG slices. **(J)** Confusion matrix of the neural network classifying interactions between tumor cells and the microenvironment.

**Supplementary figure 4. Glioblastoma tumor cells activate microglia at tumor progressing edge in perivascular invading solid tumors**. The whole tumors from **(A)** U3013MG **(B)** U3054MG **(C)** U3180 slice culture at the end point of time-lapse imaging stained with human cell marker STEM121 (green), vessels (lectin 594, red), and microglia activation marker Iba1 (yellow). The core and edge representative sites are depicted in dashed line squares for zoom-in snippets. Arrowheads indicate tumor cells interacting with activated microglia at the same location. Scale bar: 100 µm (zoomed).

**Supplementary figure 5. Treatment efficacies of dasatinib and thapsigargin on tumor cell migration and invasion in PDX slices. (A - D)** Representative first and last time point images of U3180MG, U3220MG, U3054MG, and U3179MG slice cultures untreated or treated with Dasatinib (32 μM) and Thapsigargin (4 μM). Vessels (red), tumor cells (green). Dotted line represents the tumor border at start of time lapse imaging. Scale bar: 100 μm. (E) Quantification of dose dependent effect of dasatinib and thapsigargin on cell velocity in U3013MG slices. (F) Heatmaps of the dose-dependent effects that dasatinib and thapsigargin exert on five measures of movement and growth, for all PDCX models respectively. Values are normalized estimates from fitting an LME for each model. (G-H) Differential state switching hierarchies of indicated PDCX models treated with (G) dasatinib, and (H) thapsigargin. Red and blue edges indicate an upregulation and downregulation, respectively, of the transition rates relative to control. Edge thickness represent the relative frequency of transition. (I) Distributions of morphological states for U3179MG model, showing the effect of dasatinib and thapsigargin treatment on fractions to control.

**Supplementary figure 6. Post time-lapse immunolabeling reveals distinct drug-induced effects between PDX slices. (A)** Representative snippets from the tumor core of U3013MG slices showing dose dependent effect of dasatinib on cell proliferation and resistance to thapsigargin. Scale bar: 100 µm. **(B)** Representative images from the tumor core of U3054MG slices demonstrating high Ki67 signals and resistance to thapsigargin.

## Notes

### Competing Interest Statement

The authors have declared no competing interest.

## References

1. Baker, G.J., et al., Mechanisms of glioma formation: iterative perivascular glioma growth and invasion leads to tumor progression, VEGF-independent vascularization, and resistance to antiangiogenic therapy. Neoplasia, 2014. 16(7): p. 543–61.

2. Stupp, R., et al., Radiotherapy plus concomitant and adjuvant temozolomide for glioblastoma. N Engl J Med, 2005. 352(10): p. 987–96.

3. Salvalaggio, A., et al., White Matter Tract Density Index Prediction Model of Overall Survival in Glioblastoma. Jama Neurology, 2023. 80(11): p. 1222–1231.

4. Curtin, L., et al., Shape matters: morphological metrics of glioblastoma imaging abnormalities as biomarkers of prognosis. Scientific Reports, 2021. 11(1).

5. Sorensen, A.G., et al., A “Vascular Normalization Index” as Potential Mechanistic Biomarker to Predict Survival after a Single Dose of Cediranib in Recurrent Glioblastoma Patients. Cancer Research, 2009. 69(13): p. 5296–5300.

6. Mandel, J.J., et al., Leptomeningeal dissemination in glioblastoma; an inspection of risk factors, treatment, and outcomes at a single institution. Journal of Neuro-Oncology, 2014. 120(3): p. 597–605.

7. Drumm, M., et al., Extensive brainstem infiltration is a common feature of end-stage cerebral glioblastomas. Journal of Neuropathology and Experimental Neurology, 2019. 78(6): p. 526–526.

8. Barker, F.G., et al., Survival and functional status after resection of recurrent glioblastoma multiforme. Neurosurgery, 1998. 42(4): p. 709–720.

9. Brown, T.J., et al., Association of the Extent of Resection With Survival in Glioblastoma A Systematic Review and Meta-analysis. Jama Oncology, 2016. 2(11): p. 1460–1469.

10. Al-Dalahmah, O., et al., Re-convolving the compositional landscape of primary and recurrent glioblastoma reveals prognostic and targetable tissue states. Nature Communications, 2023. 14(1).

11. Seano, G. and R.K. Jain, Vessel co-option in glioblastoma: emerging insights and opportunities. Angiogenesis, 2020. 23(1): p. 9–16.

12. Krusche, B., et al., EphrinB2 drives perivascular invasion and proliferation of glioblastoma stem-like cells. Elife, 2016. 5.

13. Pichol-Thievend, C., et al., VC-resist glioblastoma cell state: vessel co-option as a key driver of chemoradiation resistance. Nature Communications, 2024. 15(1).

14. Brooks, L.J., et al., The white matter is a pro-differentiative niche for glioblastoma. Nat Commun, 2021. 12(1): p. 2184.

15. Cuddapah, V.A., et al., A neurocentric perspective on glioma invasion. Nat Rev Neurosci, 2014. 15(7): p. 455–65.

16. Gillespie, S. and M. Monje, An active role for neurons in glioma progression: making sense of Scherer’s structures. Neuro Oncol, 2018. 20(10): p. 1292–1299.

17. Barron, T., et al., GABAergic neuron-to-glioma synapses in diffuse midline gliomas. Nature, 2025.

18. Patel, A.P., et al., Single-cell RNA-seq highlights intratumoral heterogeneity in primary glioblastoma. Science, 2014. 344(6190): p. 1396–1401.

19. Neftel, C., et al., An Integrative Model of Cellular States, Plasticity, and Genetics for Glioblastoma. Cell, 2019. 178(4): p. 835–849 e21.

20. Schmitt, M.J., et al., Phenotypic Mapping of Pathologic Cross-Talk between Glioblastoma and Innate Immune Cells by Synthetic Genetic Tracing. Cancer Discovery, 2021. 11(3): p. 754–777.

21. Larsson, I., et al., Reconstructing the regulatory programs underlying the phenotypic plasticity of neural cancers. Nat Commun, 2024. 15(1): p. 9699.

22. Larsson, I., et al., Modeling glioblastoma heterogeneity as a dynamic network of cell states. Molecular Systems Biology, 2021. 17(9).

23. Gerlee, P. and S. Nelander, The impact of phenotypic switching on glioblastoma growth and invasion. PLoS Comput Biol, 2012. 8(6): p. e1002556.

24. Farin, A., et al., Transplanted glioma cells migrate and proliferate on host brain vasculature: a dynamic analysis. Glia, 2006. 53(8): p. 799–808.

25. Dirkse, A., et al., Stem cell-associated heterogeneity in Glioblastoma results from intrinsic tumor plasticity shaped by the microenvironment. Nat Commun, 2019. 10(1): p. 1787.

26. Marques-Torrejon, M.A., E. Gangoso, and S.M. Pollard, Modelling glioblastoma tumour-host cell interactions using adult brain organotypic slice co-culture. Disease Models & Mechanisms, 2018. 11(2).

27. Minami, N., et al., Organotypic brain explant culture as a drug evaluation system for malignant brain tumors. Cancer Medicine, 2017. 6(11): p. 2635–2645.

28. Ravi, V.M., et al., Human organotypic brain slice culture: a novel framework for environmental research in neuro-oncology. Life Sci Alliance, 2019. 2(4).

29. Merz, F., et al., Organotypic slice cultures of human glioblastoma reveal different susceptibilities to treatments. Neuro-Oncology, 2013. 15(6): p. 670–681.

30. Henrik Heiland, D., et al., Tumor-associated reactive astrocytes aid the evolution of immunosuppressive environment in glioblastoma. Nat Commun, 2019. 10(1): p. 2541.

31. Griveau, A., et al., A Glial Signature and Wnt7 Signaling Regulate Glioma-Vascular Interactions and Tumor Microenvironment. Cancer Cell, 2018. 33(5): p. 874–889 e7.

32. Doroszko, M., et al., The invasion phenotypes of glioblastoma depend on plastic and reprogrammable cell states. bioRxiv, 2024: p. 2024.04.23.589925.

33. Venkataramani, V., et al., Glioblastoma hijacks neuronal mechanisms for brain invasion. Cell, 2022. 185(16): p. 2899-+.

34. Schubert, M.C., et al., Deep intravital brain tumor imaging enabled by tailored three-photon microscopy and analysis. Nat Commun, 2024. 15(1): p. 7383.

35. Lu-Emerson, C., et al., Lessons from anti-vascular endothelial growth factor and anti-vascular endothelial growth factor receptor trials in patients with glioblastoma. J Clin Oncol, 2015. 33(10): p. 1197–213.

36. Watkins, S., et al., Disruption of astrocyte-vascular coupling and the blood-brain barrier by invading glioma cells. Nat Commun, 2014. 5: p. 4196.

37. Villacampa, N., et al., Tomato lectin histochemistry for microglial visualization. Methods Mol Biol, 2013. 1041: p. 261–79.

38. Rosen, E., et al., Inference of glioblastoma migration and proliferation rates using single time-point images. Commun Biol, 2023. 6(1): p. 402.

39. Alieva, M., et al., Intravital imaging of glioma border morphology reveals distinctive cellular dynamics and contribution to tumor cell invasion. Sci Rep, 2019. 9(1): p. 2054.

40. Comba, A., et al., Spatiotemporal analysis of glioma heterogeneity reveals COL1A1 as an actionable target to disrupt tumor progression. Nature Communications, 2022. 13(1).

41. Eddy, C.Z., et al., Morphodynamics facilitate cancer cells to navigate 3D extracellular matrix. Scientific Reports, 2021. 11(1).

42. Xie, Y., et al., The Human Glioblastoma Cell Culture Resource: Validated Cell Models Representing All Molecular Subtypes. Ebiomedicine, 2015. 2(10): p. 1351–1363.

43. Johansson, P., et al., A Patient-Derived Cell Atlas Informs Precision Targeting of Glioblastoma. Cell Reports, 2020. 32(2).

44. Almstedt, E., et al., Real-time evaluation of glioblastoma growth in patient-specific zebrafish xenografts. Neuro-Oncology, 2022. 24(5): p. 726–738.

45. Lee, K.Y., et al., Video Stabilization using Robust Feature Trajectories. 2009 Ieee 12th International Conference on Computer Vision (Iccv), 2009: p. 1397–1404.

46. Guizar-Sicairos, M., S.T. Thurman, and J.R. Fienup, Efficient subpixel image registration algorithms. Opt Lett, 2008. 33(2): p. 156–8.

